# Identification of Entomopathogenic Nematode isolates from Nigeria and their pathogenicity against the invasive fall armyworm, *Spodoptera frugiperda* J.E. Smith (Lepidoptera: Noctuidae)

**DOI:** 10.1101/2025.11.23.689995

**Authors:** CT. Okolo, A. Claudius-Cole, FW. Grundler, C. Borgemeister

**Affiliations:** University of Bonn, Center for Development Research (ZEF), Germany; University of Ibadan, Crop Protection and Environmental Biology, Nigeria; University of Bonn, Institute of Crop Science and Resource Conservation (INRES).; Ahmadu Bello University, Department of Crop Protection, Nigeria

## Abstract

The fall armyworm (FAW, *Spodoptera frugiperda* J.E. Smith [Lep.: Noctuidae]), a highly destructive lepidopteran pest, poses a serious threat to maize and other staple crops across sub-Saharan Africa. With growing concerns over resistance to synthetic insecticides, entomopathogenic nematodes (EPNs) have emerged as viable biological control alternative. This study reports the isolation, identification, and virulence evaluation of six indigenous EPN isolates collected from different agroecological zones in Nigeria. Morphological and morphometric assessments, supported by molecular analyses of internal transcribed spacer regions (ITS), D2–D3, and mitochondrial gene regions, identified the isolates as *Heterorhabditis bacteriophora*, *Steinernema carpocapsae*, *S. feltiae*, *S. nepalense*, and *Oscheius myriophilus*. Virulence bioassays were conducted against four developmental stages of *S. frugiperda* (2^nd^, 4^th^, 6^th^ instar larvae and pupae) using four dosages (25, 50, 100, and 200 infective juveniles [IJs]/insect). All isolates exhibited dose- and time-dependent pathogenicity, with FAW’s 2^nd^ instar larvae showing the highest susceptibility. *Heterorhabditis bacteriophora* (Ib-CRIN68) caused the highest mortality (82.4 ± 5.6%) and the lowest lethal concentration (LC₅₀) and lethal time (LT₅₀) values among the tested EPN isolates. Significant effects of isolate, dosage, and exposure time on larval mortality (p < 0.001) were found, particularly in early instars. The findings highlight the high biocontrol potential of these Nigerian EPN isolates and underscore the value of integrating molecular and morphological diagnostics for accurate species identification. This work provides a critical foundation for the future ecological characterization and field deployment of indigenous nematodes in sustainable pest management strategies against *S. frugiperda* in West Africa.

## Introduction

Fall armyworm (FAW), *Spodoptera frugiperda* J.E. Smith (Lep.: Noctuidae), has rapidly emerged as a severe invasive pest in Africa, significantly impacting maize productivity and food security globally, particularly in sub-Saharan Africa (SSA) (Kenis et al., 2023). Native to tropical and subtropical regions of the Americas, FAW was first reported in West and Central Africa in 2016, quickly spreading across the continent and causing considerable economic losses (Agboyi et al., 2020; Goergen, Kumar, Sankung, Togola, & Tamò, 2016). Its polyphagous nature, rapid development, and high reproductive capacity enable it to infest numerous economically important crops, especially maize, resulting in yield losses ranging from 20 to 50% in severe cases (Kenis et al., 2023; Overton et al., 2021). The current management of FAW in SSA relies heavily on synthetic insecticides; however, this approach is increasingly associated with environmental hazards, development of resistance, and human health concerns, emphasizing the need for safer and more sustainable alternatives (De Groote et al., 2020; Hruska, 2019; Kenis et al., 2023; Makale et al., 2022; Odong et al., 2024).

Entomopathogenic nematodes (EPNs), belonging primarily to the genera *Steinernema* and *Heterorhabditis*, are prominent biological control agents used globally against diverse insect pests due to their effectiveness, rapid killing action, host-specificity, environmental safety, and compatibility with integrated pest management (IPM) programmes (Ehlers, 2007; Machado et al., 2025; Šreibr et al., 2025). Despite the proven potential of EPNs, their application in SSA, particularly Nigeria, remains limited primarily due to inadequate characterization and documentation of indigenous species (Akyazi, Ansari, Ahmed, Crow, & Mekete, 2012; Daramola et al., 2021). Indigenous nematode isolates often exhibit superior adaptation and efficacy against local pest populations compared to exotic isolates, reinforcing the importance of isolating, identifying, and characterizing local EPN species for effective pest management (Dichusa, Ramos, Aryal, Sumaya, & Sumaya, 2021; Guide et al., 2024).

Accurate identification of EPN isolates, using both morphological and molecular techniques, is crucial for their effective use and commercialization in biological control programs. Morphological characterization, including detailed morphometric analyses, provides initial identification; however, molecular approaches, such as the sequencing of internal transcribed spacer (ITS) regions and mitochondrial DNA markers, offer more precise species identification, and enabling the determination of phylogenetic relationships and genetic variability (Aryal et al., 2021; Gumussoy et al., 2022). Such detailed identification facilitates the selection of highly virulent and ecologically adapted isolates suitable for local pest management strategies.

This study addresses the knowledge gap in the characterization and bio-efficacy of indigenous Nigerian EPN isolates against FAW. We isolated and identified local EPN isolates from Ibadan and Zaria of south-western and north-western Nigeria, respectively, using morphological and molecular tools, assessed their virulence against different developmental stages of FAW, and tested their potential as more sustainable biological control agents within an IPM framework. Understanding the effectiveness and adaptability of these indigenous EPN isolates not only aids in local pest management but also contributes to the global pool of biologically effective agents, promoting sustainable agriculture and food security in SSA.

## Materials and Methods

### Study Area and Sampling Design

The study was conducted across two distinct agroecological zones in Nigeria: Ibadan in the southwestern region and Zaria in the northwestern region.

In Ibadan, a major West African city located in Oyo State within the lowland rainforest agroecological zone, characterized by humid tropical climate conditions, soil samples were collected from seven different locations. The coordinates of these sampling sites were as follows: Moniya (7.502261, 3.909373), Ajibode (7.456701, 3.884341), Idi Ayunre-CRIN (7.235996, 3.866177), Idi Ishin-FRIN (7.391910, 3.863259), Oganla (7.404751, 3.846373), UI-CPEB (7.450322, 3.896921) and Apata (7.386540, 3.842505). In total, 128 soil samples were collected from these sites in and around Ibadan.

The city of Zaria, situated in Kaduna State within the Northern Guinea savannah agroecological zone, characterized by a relatively dry climate, grassy vegetation, and moderate rainfall patterns, served as the second study area. Soil samples were collected from five distinct locations in and around Zaria, represented by the coordinates: ABU Dam (11.133388, 7.654355), Samaru (11.165714, 7.633078), Shika (11.204446, 7.560817), Hanwa (11.133287, 7.712618), and Chikaji (11.129026, 7.713202). A total of 74 soil samples were collected from these locations.

The soil sampling procedure involved the random collection of approximately 1.5 kg of soil from the upper 0–15 cm soil layer at each sampling location. The samples were placed into labelled polyethylene bags, sealed, and promptly transported to the laboratory to ensure minimal disturbance to the sample’s integrity.

Isolation of EPNs was then performed. Prior to nematode isolation, a representative portion of each soil sample was analysed to determine soil composition parameters, including soil type, texture, pH, moisture content, and organic matter content. For the isolation process, each soil sample was individually processed using the insect bait method with larvae of the wax moth *Galleria mellonella* (L.) (Lep.: Pyralidae). Approximately 1 kg of soil from each sample was placed in lid-covered plastic boxes and clearly labelled. Ten final-instar larvae of *G. mellonella* were placed on the surface of each soil sample and incubated in the dark under controlled conditions of 25°C temperature and 55% relative humidity (r.h.), following the methodology of Bedding and Akhurst (1975).

Larvae were monitored every 24 hours for up to seven days until complete mortality was observed. Dead larvae exhibiting typical nematode infection symptoms were subsequently transferred to modified White traps (White, 1927) to collect emerging infective juveniles (IJs) of the EPNs. The pathogenicity of the recovered nematode isolates was confirmed by reinfestation tests using fresh *G. mellonella* larvae. Newly emerged IJs collected from the White traps were washed with sterile distilled water and subsequently stored at 13°C for further characterization and bioassays. From all the sampled locations, a total of six EPN isolates were successfully recovered, five isolates originating from Ibadan and one from Zaria.

### Morphological and Morphometric Identification

Morphological and morphometric characterization of each EPN isolate was conducted by examining IJs, males, females, and hermaphrodites. Specifically, 20 specimens from each life stage were randomly selected for detailed measurements. Specimens were identified at the genus and species level using a standard taxonomic key (Nguyen & Smart, 1996), specifically designed for distinguishing EPNs belonging to families Steinernematidae and Heterorhabditidae.

All morphometric measurements (expressed in micrometers, µm) were obtained using a phase-contrast microscope (Nikon Eclipse 80i) equipped with the Nikon DS-L2 image acquisition software, ensuring high accuracy and consistency in measurements.

### Molecular Characterization

#### DNA Extraction

For DNA extraction, genomic DNA from individual specimens of each of the six nematode isolates was extracted using the ROTI Prep Genomic DNA Mini 2.0 Extraction Kit (Carl Roth GmbH). Single virgin females were initially washed separately in Ringer’s solution and subsequently rinsed with phosphate-buffered saline (PBS, pH 7.2). Each specimen was then individually transferred into sterile polymerase chain reaction (PCR) tubes (0.2 mL) containing 20 μL of extraction buffer composed of 17.6 μL nuclease-free distilled water, 2 μL of 5× PCR buffer, 0.2 μL of 1% Tween, and 0.2 μL of proteinase K. The samples were either frozen at −20°C for 60 minutes or overnight, then immediately incubated in a PCR thermocycler at 65 °C for 1.2 hours followed by incubation at 95 °C for 10 minutes. Subsequently, the lysates were cooled on ice, centrifuged at 6,500 × g for 3 minutes, and the resulting supernatants were stored at −20 °C and used as DNA templates for PCR amplification.

#### Amplification of Target Gene Regions

Four taxonomically relevant gene regions were targeted for amplification using the PCR technique. The internal transcribed spacer (ITS1-5.8S-ITS2) regions were amplified using specific primers: the forward primer (5′-TGATTACGTCCCTGCCCTTT-3′) and the reverse primer (5′-TTTCACTCGCCGTTACTAAGG-3′). Similarly, the D2–D3 expansion segments of the 28S rRNA were amplified with primers D2F (forward: 5′-CCTTAGTAACGGCGAGTGAAA-3′) and 536 (reverse: 5′-CAGCTATCCTGAGGAAAC-3′). For the 12S mitochondrial gene, primers 505F (forward: 5′-GTTCCAGAATAATCGGCTAGAC-3′) and 506 R (reverse: 5′TCTACTTTACTACAACTTACTCCCC-3′) were employed. Lastly, amplification of the mitochondrial Cytochrome Oxidase subunit I (MT-COI) gene was achieved using primers HCF (forward: 5′-TTACATGATACTTATTATG-3′) and HCR (reverse: 5′-CTGATAACTGTGACCAAATACATA-3′).

The PCR reactions were conducted in 25 µL volumes comprising 2 µL DNA extract, 12.5 µL DreamTaq Green PCR Master Mix (Thermo Scientific, USA), 0.75 µL of each forward and reverse primer (10 µM each), and 9 µL nuclease-free distilled water. Amplifications were performed in an Applied Biosystems Veriti 96-Well Thermal Cycler (Thermo Scientific, USA) under the following thermal cycling conditions: for ITS, D2–D3, and 12S genes, initial denaturation at 94 °C for 3 minutes was followed by 35 cycles of denaturation at 94 °C for 30 seconds, annealing at 50 °C for 30 seconds, extension at 72 °C for 1 minute 30 seconds, and a final elongation step at 72 °C for 20 minutes. For the MT-COI gene, initial denaturation at 94 °C for 3 minutes was followed by 38 cycles of denaturation at 94 °C for 10 seconds, annealing at 40 °C for 30 seconds, extension at 72 °C for 60 seconds, and a final elongation step at 72 °C for 10 minutes. The PCR products (5 μL each) were separated by electrophoresis on a 1% agarose gel buffered with Tris–boric acid–EDTA (TBE) and stained with SYBR Safe DNA Gel Stain Invitrogen, Carlsbad, CA, (Thermo Scientific, USA) for visualization after electrophoresis at 100 V for 45 minutes.

Sequencing of the PCR products was conducted using the Qiagen QIAquick Gel Extraction Kit, with bidirectional sequencing (using both forward and reverse primers) (Eurofins Genomics, Germany). Phylogenetic analyses were completed by retrieving genomic sequences corresponding to all valid described species of the genera *Heterorhabditis*, *Steinernema*, and *Oscheius* from the National Center for Biotechnology Information (NCBI) database using the Basic Local Alignment Search Tool (BLAST). Sequence alignments were performed using MUSCLE (v3.8.31), and phylogenetic relationships were reconstructed using the maximum likelihood (ML) method in MEGA 11 software, based on nucleotide substitution models selected through best-fit substitution model analysis. The models applied were the Tamura–Nei model (TN93+G+I) for MT-COI sequences and the Kimura 2-parameter model (K2+G) for D2–D3 and ITS sequences. ML phylogenetic trees were constructed from initial trees derived from neighbour-joining (NJ) and BioNJ algorithms, based on matrices of pairwise distances estimated by maximum composite likelihood methods. Bootstrap support values were indicated on tree branches, with parameters of discrete gamma distribution (+G) and evolutionary invariable sites (+I) applied where necessary. Trees were drawn to scale, with branch lengths measured in substitutions per site.

### Insect Rearing and Preparation for Virulence Bioassays

FAW larvae were maintained and reared under controlled quarantine conditions at the Insect Rearing Facility of the Entomology Unit at the International Institute for Tropical Agriculture (IITA) in Ibadan, Nigeria. Insects originated from a laboratory colony of *S. frugiperda*, established from specimens collected from a maize field in the vicinity of Ibadan, and maintained in a controlled environment (25 ± 1°C, 50 ± 5% r.h.). FAW larvae were nurtured on an artificially formulated maize-based diet, specifically designed for lepidopteran caterpillars (Chen, Mhlungu, Chang, & Kafle, 2023; Greene, Leppla, & Dickerson, 1976). Adult insects and the 1^st^ to 3^rd^ instar larvae were cultured in large round plastic containers (diameter of 30 cm) covered with netting (40-mesh). The 4^th^ to 6^th^ instar larvae were individually cultured in plastic cups (diameter of 4.5 cm) that had eight small holes (1 mm diameter) in the lid.

The insects were segregated based on their developmental stages (2^nd^, 4^th^, 6^th^ instars, and pupae) and kept in separate culture plates to prevent mixing and cannibalism. Before starting the bioassays, insects were acclimated to ambient laboratory conditions (approximately 25 ± 2°C room temperature and 55 ± 10% r.h.) for 24 hours, ensuring uniform physiological conditions for the experiments.

The larvae and pupae were then used in controlled virulence bioassays to assess their susceptibility to the isolated EPN strains. Similar-sized 2^nd^, 4^th^, and 6^th^ instar larvae, as well as pupae, were selected for the bioassays.

### Virulence Bioassays

Virulence bioassays were conducted to evaluate the susceptibility of FAW to six indigenous EPN isolates. The nematode isolates tested were Ib-CRIN68, Ib-IART45, Ib-ITUC102, Ib-FRIN32, and Ib-HORT from Ibadan, and Za-SAM from Zaria. Tests were carried out with 2^nd^, 4^th^, and 6^th^ instar larvae and newly formed FAW pupae. Before the bioassays, insects were acclimated at room temperature (approximately 25 ± 2 °C, 55 ± 10% r.h.) for 24 hours to ensure consistent physiological conditions.

Bioassays were set up in sterile 90 mm Petri dishes, each lined with qualitative filter paper. For assays involving larval stages, approximately 2.5 g of the artificial diet was evenly distributed over the filter paper in each dish, and five larvae of the same developmental stage (2^nd^, 4^th^, or 6^th^ instar) were gently transferred onto the diet surface. For pupal bioassays, dishes were prepared with moistened filter paper only, onto which five pupae were placed carefully without diet.

To prepare the nematode suspensions, IJs of each isolate were cultured, harvested, and accurately counted under a stereomicroscope. Four different dosages of nematodes (25, 50, 100, and 200 IJs per insect) were prepared in sterile distilled water. Specifically, for each Petri dish containing five insects, suspensions contained a total of 125, 250, 500, or 1,000 IJs in 1 mL of distilled water, respectively.

Nematode suspensions were applied uniformly onto the filter paper surface using a micropipette, carefully ensuring even distribution without disturbing the insects. Control groups for each FAW developmental stage were treated similarly but received only sterile distilled water. Petri dishes were then tightly sealed with Parafilm to maintain constant humidity levels and incubated in a controlled environment chamber at 25 ± 1 °C and 60 ± 5% r.h.

Mortality was monitored at 24-, 48-, and 72-hours post-application. During each observation, larvae were gently prodded under a stereomicroscope, and lack of movement or response was recorded as mortality. Pupae were visually examined for changes in coloration or signs of abnormal development as an indication of mortality. Data on insect survival and mortality were carefully recorded for each treatment at each observation period. To confirm nematode-induced mortality, a subset of insect cadavers was dissected and examined microscopically for the presence of EPNs.

Each combination of nematode isolate, dosage, and FAW developmental stage was tested with a total of ten insects, divided into two replicate dishes (five insects per dish). The entire bioassay experiment was independently repeated three times on separate dates to ensure robustness and reproducibility of the results.

### Data Analysis

Mortality data were subjected to statistical analyses, including analysis of variance (ANOVA) or logistic regression, to examine the potential differences among isolates, dosages, and FAW stages. Post-hoc comparisons were conducted using Tukey’s HSD test to identify significant differences between treatment means. Additionally, dose–response curves depicting percent mortality relative to nematode dosage were generated for each isolate and developmental stage.

## Results

### Nematode Isolation

A total of 202 soil samples were collected from twelve locations in the two agroecological zones of Nigeria, i.e., Ibadan (128 samples) in the southwest rainforest (tropic warm subhumid) and Zaria (74 samples) in the northern savannah (tropic warm semiarid) (Fig. 1). Five EPN isolates were found in Ibadan, with positive recoveries at Moniya (5.0%), Ajibode (5.6%), Idi Ayunre-CRIN (4.2%), Idi Ishin-FRIN (6.2%), and Oganla (5.6%). No isolates were recovered from UI-CPEB and Apata. In Zaria, a single isolate was recovered at Samaru (7.1%), but none from ABU Dam, Shika, Hanwa, or Chikaji. The overall recovery rates were 3.9% for Ibadan and 1.3% for Zaria (Fig. 2).

**Fig. 1.**
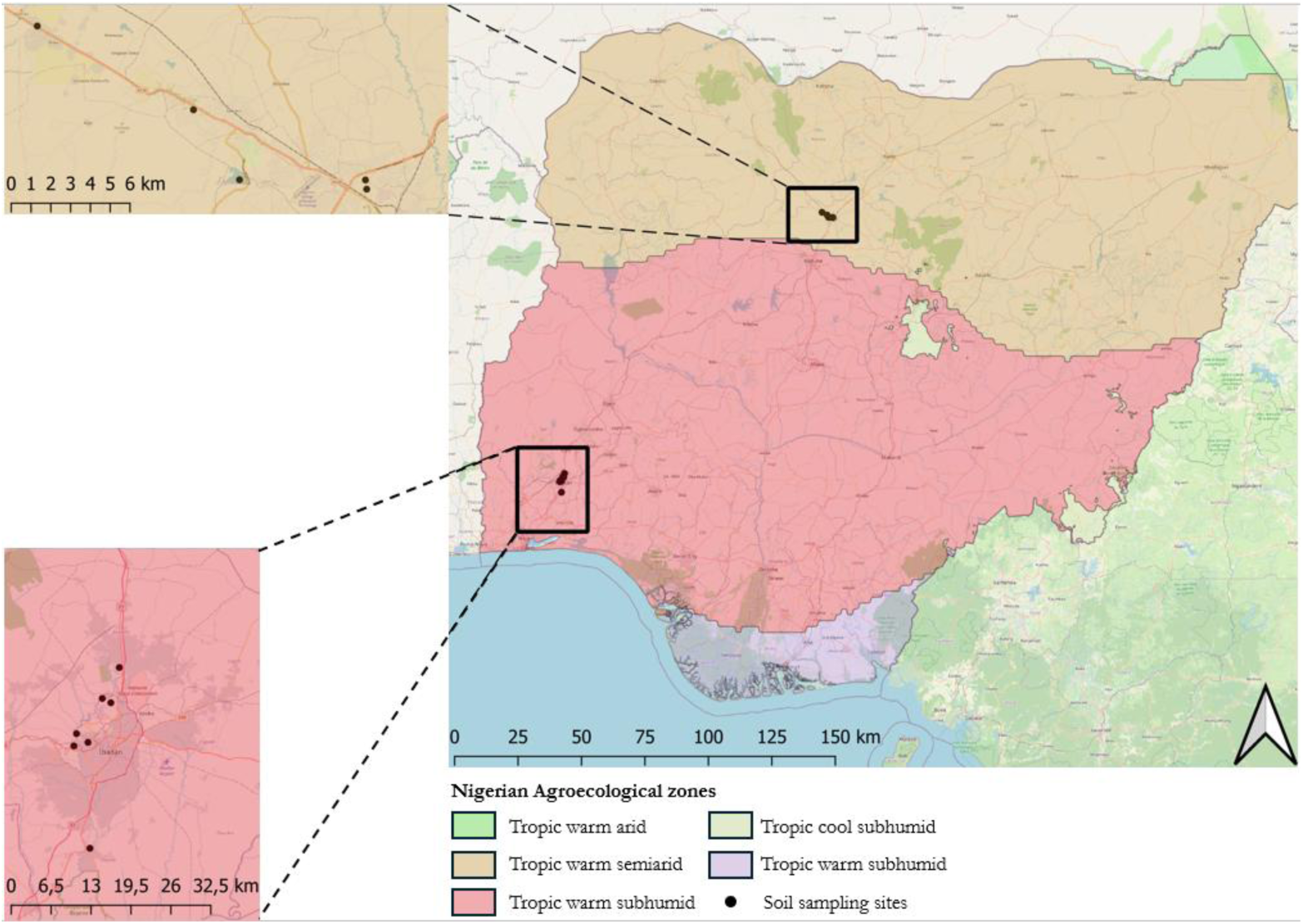
Map of Nigeria showing the agroecological zones and soil sampling sites for entomopathogenic nematode (EPN) isolation, with insets highlighting the specific sampling locations in each region.

**Fig. 2.**
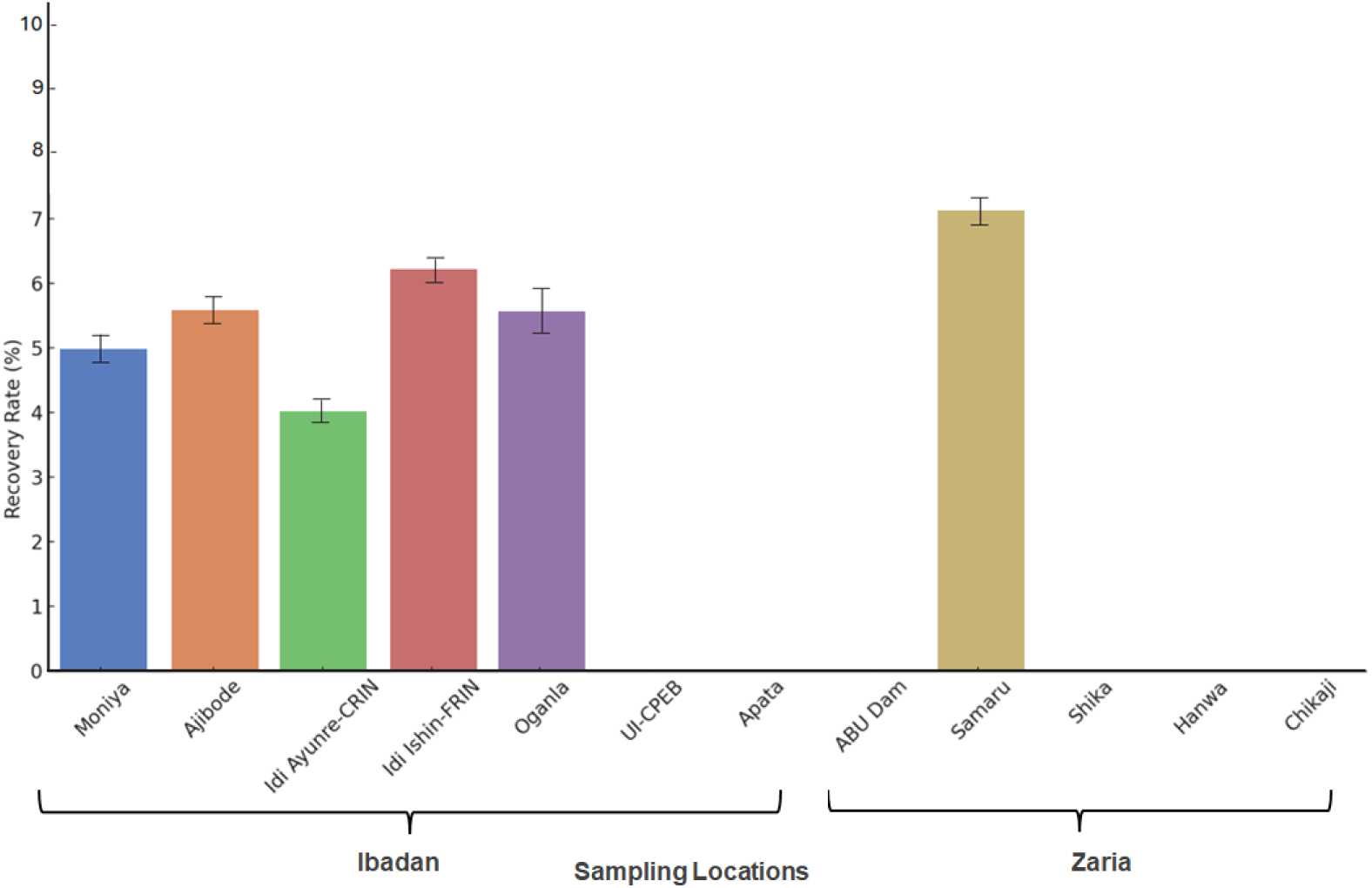
Recovery rate of entomopathogenic nematode (EPN) isolates from soil samples collected across twelve locations in Nigeria. Bars represent the percentage of soil samples from each site yielding at least one isolate.

### Morphological and morphometric Identification

Morphological characterization of the six recovered EPN isolates was performed based on detailed morphometric analyses of IJs, males, amphimictic females, and hermaphrodites, averaging measurements from 20 specimens per life stage for each isolate Examination focused on key morphological characters including body length (L), maximum body diameter (MBD), tail length (T), anal body diameter (ABD), position of the excretory pore (EP), nerve ring distance (NR), pharynx length (ES), and calculated morphometric ratios (a, b, c, D%, and E%)(Fig. 3).

**Fig. 3.**
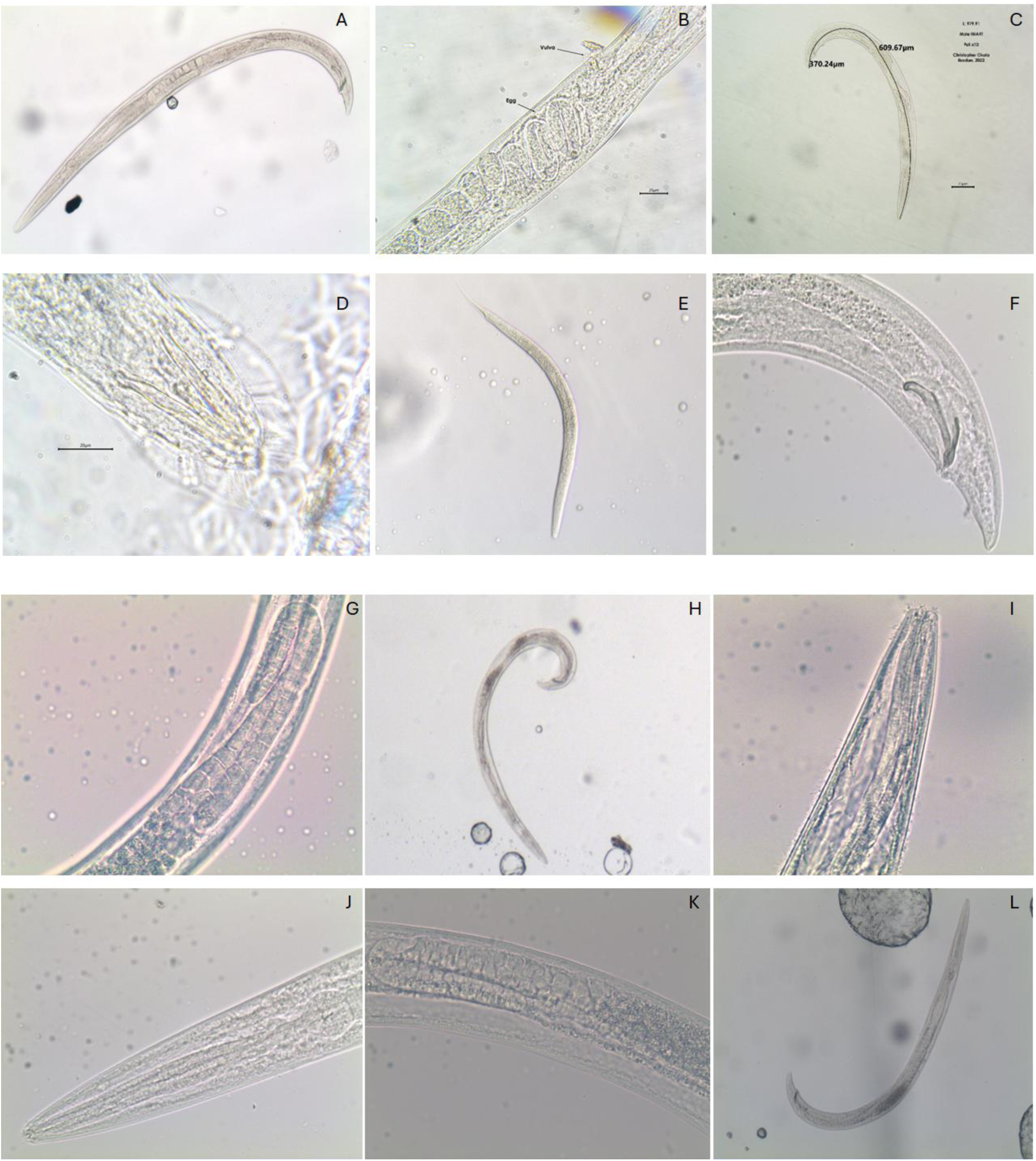
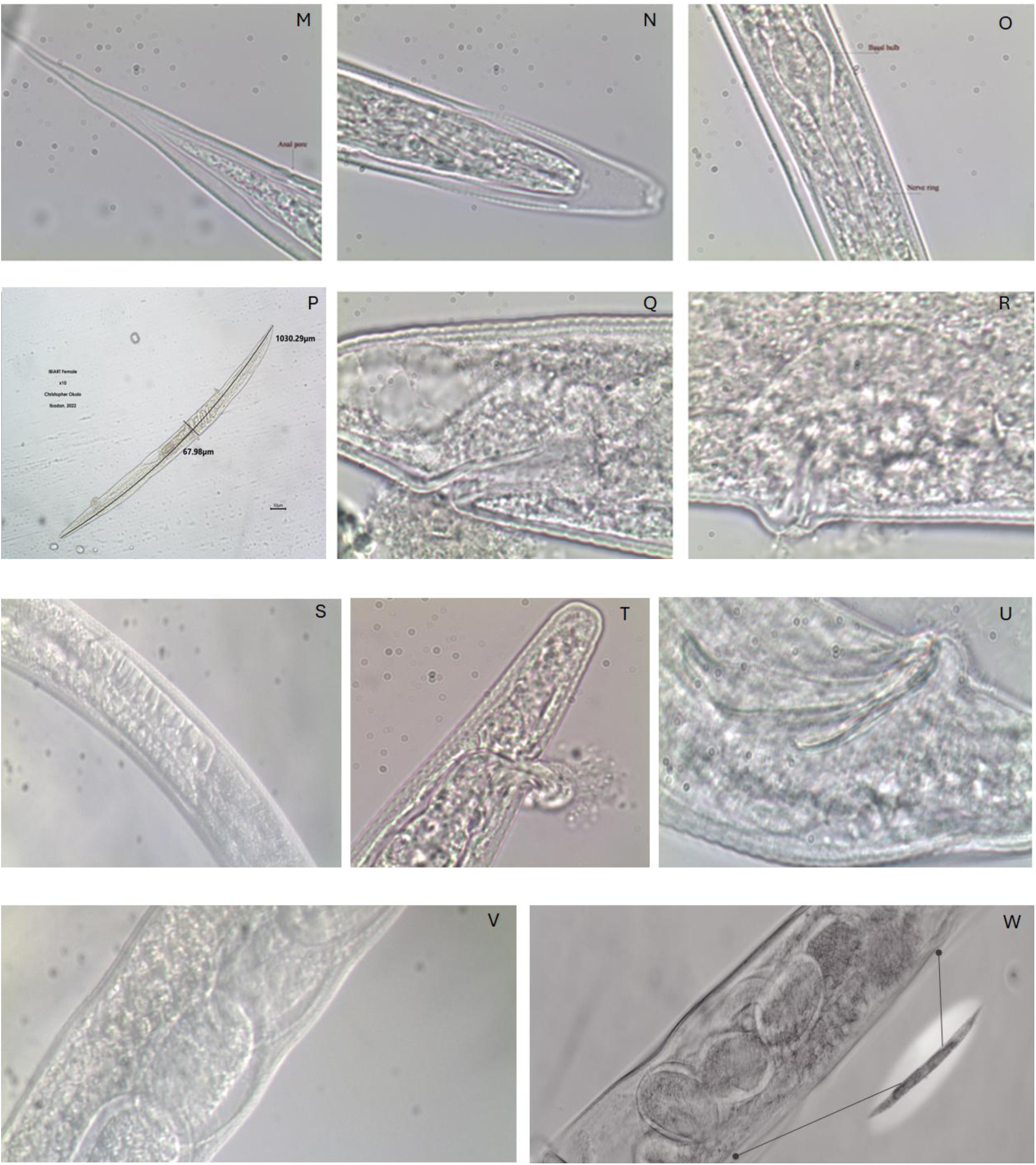
Light micrographs of different life stages of entomopathogenic nematodes (EPN) isolated from Nigeria. A: Full length male sample of Ib-ITUC102, B: Ib-IART45 female showing matured and vulva, C: Full length male Ib-IART45, D: Tail region of Ib-CRIN68 showing the distinct bursar of *Heterorhabditis* sp., E: Infective juvenile (IJ) of Ib-HORT, F: Tail region of Za-SAM showing the spicule and gubernaculum, G: Testis reflex of Ib-HORT, H: Male of Ib-FRIN32, I: Anterior part Ib-HORT, J: Anterior part of Ib-IART45 showing median bulb, K: Testis reflex of IB-ITUC102, L: Full length male of Za-SAM, M: Tail region of Ib-ITUC102 IJ showing hyalin, N: Head region of Ib-ITUC102 juvenile showing outer sheath. O: Median and nerve ring of Ib-FRIN32, P: Full length of female of Ib-IART45, Q: Anal region of Za-SAM showing excrete, R: Vulva region of ZA-SAM, S: Fat deposit in Ib-CRIN68 IJ, T: Tail region of Ib-FRIN32 showing excretes, U: Spicule and gubernaculum of IB-IART45 male, V: Eggs in female IB-CRIN68, W: Eggs in female Ib-FRIN32.

Morphometric analysis revealed distinct genus- and isolate-level traits across the three examined life stages. In the male stage (Table 1), Ib-CRIN68 exhibited typical *Heterorhabditis* morphology, with elongate body form (∼1,100 µm), strongly curved testis reflexion, and well-developed spicules (∼50 µm). Males of -FRIN32 were notably shorter (∼780 µm) but had prominently curved testes and longer spicules (∼42 µm), consistent with *Oscheius* diagnostics. *Steinernema* isolates showed moderate variation in male body size and tail morphometrics, with Ib-IART45, Ib-ITUC102 showing body lengths of ∼1,000 µm and high spicule-to-body diameter ratios, while Za-SAM and Ib-HORT presented longer males (up to 1,000 µm). In the female stage (Table 2), females of Ib-CRIN68 were the longest among all isolates (∼1,350 µm), with clearly defined vulval positioning and a long tail (∼120 µm). Ib-HORT followed similar patterns with large body size (∼1,200 µm) and cruiser-like tail characteristics. Other *Steinernema* isolates had more compact females, while Ib-FRIN32 showed distinctly smaller females (∼900 µm) and reduced maximum body diameter, differentiating it clearly at the genus level. In the IJs stage (Table 3), all *Steinernema* isolates displayed morphometrics typical of ambushers and intermediates. Ib-IART45, Ib-ITUC102 IJs were shorter (∼525–540 µm) with high L/T ratios and elongated hyaline tail regions (∼45 µm), consistent with ambusher behaviour. Za-SAM exhibited intermediate IJ length (∼570 µm) and tail traits aligning with its foraging flexibility. Ib-CRIN68 had the longest IJs (∼600–630 µm), while Ib-FRIN32 IJs were more compact (∼600 µm) but had an extended hyaline region contributing nearly half the tail length, a key diagnostic trait of the genus *Oscheius*.

**Table 1.**
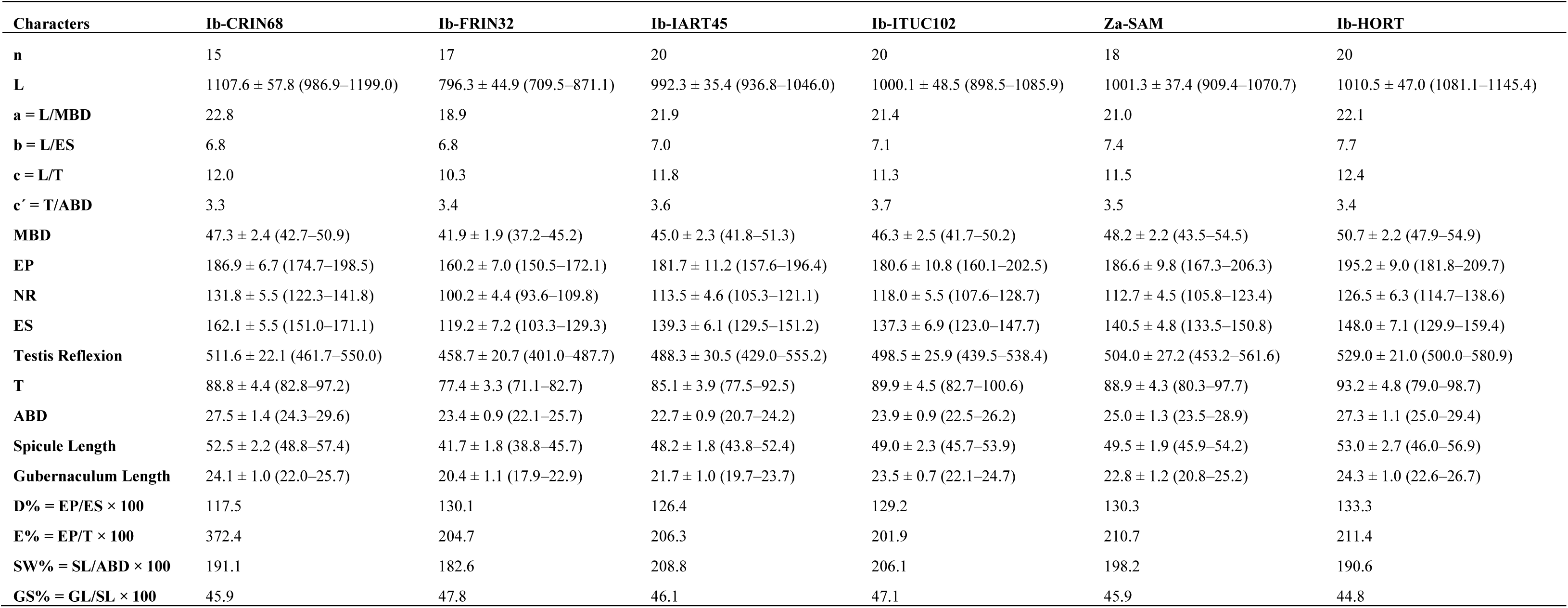
Male morphometrics of six isolates of entomopathogenic nematodes from Nigeria (all measurements are in μm except n, ratio, and percentage), and in the form: mean ± s.d. (range)

**Table 2.** Female morphometrics of six isolates of entomopathogenic nematodes from Nigeria (all measurements are in μm except n, ratio, and percentage), and in the form: mean ± s.d. (range)

| Characters | Ib-CRIN68 | Ib-FRIN32 | Ib-IART45 | Ib-ITUC102 | Za-SAM | Ib-HORT |
| --- | --- | --- | --- | --- | --- | --- |
| **n** | 17 | 18 | 20 | 19 | 20 | 18 |
| **L** | 1347.8 ± 77.3 (1214.0–1458.7) | 922.3 ± 40.8 (825.2–1007.3) | 1132.1 ± 52.6 (1038.8–1215.2) | 1114.6 ± 57.2 (995.6–1198.0) | 1263.8 ± 59.5 (1164.5–1342.0) | 1215.4 ± 71.8 (1202.0–1383.7) |
| **a = L/MBD** | 14.1 | 14.7 | 15.7 | 14.9 | 14.2 | 15.1 |
| **b = L/ES** | 7.9 | 7.2 | 7.8 | 7.9 | 8.3 | 8.6 |
| **c = L/T** | 10.7 | 9.2 | 10.2 | 10.1 | 11.0 | 10.6 |
| **c' = T/ABD** | 3.9 | 4.0 | 3.9 | 3.9 | 3.8 | 3.9 |
| **V'** | 698.7 ± 33.8 (631.5–767.0) | 491.0 ± 23.4 (447.4–539.3) | 590.7 ± 24.8 (555.6–636.6) | 606.8 ± 30.7 (545.6–678.6) | 672.9 ± 28.0 (617.6–729.4) | 712.6 ± 30.0 (674.7–786.5) |
| **V = V'/L \*100** | 51.8 | 53.2 | 52.2 | 54.4 | 53.2 | 51.9 |
| **MBD** | 94.8 ± 4.9 (85.2–103.4) | 62.7 ± 2.7 (58.1–67.1) | 71.8 ± 4.0 (64.7–78.3) | 75.2 ± 3.7 (67.6–82.4) | 87.1 ± 3.6 (80.3–93.6) | 91.5 ± 3.3 (86.4–98.5) |
| **EP** | 213.1 ± 8.1 (195.6–225.3) | 168.1 ± 8.3 (149.7–181.2) | 184.7 ± 3.9 (177.7–191.6) | 188.2 ± 7.1 (171.2–198.7) | 202.7 ± 9.5 (183.9–220.5) | 214.6 ± 9.4 (196.2–233.1) |
| **NR** | 145.4 ± 6.2 (136.2–156.7) | 110.5 ± 5.8 (96.2–121.6) | 122.6 ± 5.8 (112.2–133.1) | 126.4 ± 4.3 (118.6–133.8) | 132.8 ± 5.8 (121.9–144.8) | 140.3 ± 5.7 (127.4–148.2) |
| **ES** | 168.7 ± 9.4 (154.5–185.3) | 129.9 ± 7.5 (115.9–140.9) | 144.6 ± 6.6 (128.6–155.7) | 145.4 ± 7.0 (133.0–161.3) | 147.9 ± 6.6 (139.1–163.9) | 158.8 ± 7.2 (141.1–171.6) |
| **T** | 125.5 ± 6.8 (107.6–134.9) | 104.0 ± 4.7 (95.2–113.0) | 107.7 ± 6.5 (93.0–119.5) | 112.2 ± 6.0 (99.3–119.6) | 114.7 ± 5.4 (101.7–123.3) | 131.6 ± 7.8 (110.5–145.8) |
| **ABD** | 31.6 ± 1.5 (28.7–35.6) | 26.3 ± 1.3 (24.2–28.7) | 28.1 ± 1.6 (25.8–31.2) | 28.7 ± 1.0 (27.0–30.4) | 30.2 ± 1.2 (27.8–31.8) | 33.1 ± 1.8 (29.5–36.3) |
| **D% = EP/ES × 100** | 122.5 | 131.3 | 126.2 | 129.4 | 135.7 | 133.7 |
| **E% = EP/T × 100** | 167.9 | 166.1 | 170.1 | 169.0 | 176.0 | 165.3 |

**Table 3.** Infective juvenile morphometrics of six isolates of entomopathogenic nematodes from Nigeria (all measurements are in μm except n, ratio, and percentage), and in the form: mean ± s.d. (range)

| Characters | Ib-CRIN68 | Ib-FRIN32 | Ib-IART45 | Ib-ITUC102 | Za-SAM | Ib-HORT |
| --- | --- | --- | --- | --- | --- | --- |
| **n** | 20 | 20 | 20 | 20 | 20 | 20 |
| **L** | 569.4 ± 21.7 (530.4–615.6) | 601.9 ± 23.3 (565.2–654.8) | 536.4 ± 32.4 (474.1–590.5) | 538.0 ± 29.0 (499.0–592.9) | 570.4 ± 31.6 (514.7–617.0) | 582.0 ± 23.7 (526.4–629.9) |
| **a = L/MBD** | 21.2 | 21.5 | 22.3 | 21.7 | 23.1 | 22.4 |
| **b = L/ES** | 5.3 | 5.9 | 5.2 | 5.2 | 5.4 | 5.2 |
| **c = L/T** | 6.2 | 7.0 | 6.1 | 6.0 | 6.7 | 6.5 |
| **c' = T/ABD** | 6.0 | 6.0 | 6.2 | 6.3 | 5.6 | 6.0 |
| **MBD** | 26.3 ± 1.6 (23.4–29.5) | 28.0 ± 1.5 (25.3–31.4) | 23.9 ± 1.1 (22.2–26.0) | 25.3 ± 1.3 (22.3–27.1) | 24.9 ± 0.9 (23.1–26.4) | 26.1 ± 1.0 (24.6–27.5) |
| **EP** | 117.6 ± 7.0 (105.0–136.2) | 107.2 ± 4.8 (95.9–114.7) | 111.7 ± 5.2 (101.0–121.8) | 113.7 ± 3.9 (108.4–123.4) | 114.3 ± 4.9 (107.0–127.2) | 120.1 ± 6.0 (111.8–131.9) |
| **NR** | 86.4 ± 3.9 (79.3–91.9) | 81.6 ± 5.1 (71.2–91.1) | 86.5 ± 3.3 (77.8–92.4) | 87.8 ± 4.8 (79.2–95.9) | 84.5 ± 4.8 (77.4–91.7) | 89.9 ± 4.9 (81.4–99.1) |
| **ES** | 107.0 ± 4.0 (101.0–115.6) | 101.4 ± 6.0 (92.2–112.9) | 102.7 ± 4.5 (94.9–113.3) | 103.0 ± 5.0 (87.6–109.5) | 105.5 ± 4.0 (97.0–114.9) | 109.9 ± 4.2 (102.8–122.1) |
| **T** | 89.2 ± 5.5 (76.4–98.2) | 84.4 ± 3.2 (76.1–90.5) | 87.0 ± 4.6 (73.6–93.6) | 87.4 ± 4.9 (76.0–97.0) | 86.6 ± 3.4 (82.0–94.3) | 87.5 ± 4.6 (79.9–99.4) |
| **ABD** | 14.9 ± 0.8 (13.3–16.4) | 14.3 ± 0.7 (13.3–16.1) | 13.7 ± 0.6 (12.5–14.8) | 14.1 ± 1.0 (11.8–15.8) | 15.3 ± 0.8 (14.0–16.8) | 14.8 ± 0.8 (13.5–17.2) |
| **H** | | | 42.3 ± 1.6 (38.8–45.3) | 41.8 ± 1.9 (39.1–45.4) | 41.0 ± 2.1 (37.4–45.9) | 42.4 ± 2.9 (37.8–50.8) |
| **D% = EP/ES × 100** | 109.4 | 105.6 | 106.5 | 109.6 | 110.7 | 107.5 |
| **E% = EP/T × 100** | 130.7 | 127.2 | 125.6 | 125.6 | 134.1 | 133.8 |
| **H% = H/T × 100** |  |  | 47.6 | 48.3 | 47.5 | 47.7 |

### Molecular Identification

Molecular sequencing results and phylogenetic analyses supported and expanded upon the morphological identifications. The isolate Ib-CRIN68 demonstrated a high sequence similarity (>99%) to *H. bacteriophora*. Similarly, isolates Ib-IART45 and Ib-ITUC102 exhibited genetic congruence with *S. carpocapsae* sequences (>98% similarity), consistent with their morphological profiles. Ib-HORT was genetically closely aligned (>98% similarity) with *S. nepalense*, while Za-SAM showed high sequence similarity to *S. feltiae* (>98% identity). Molecular data for Ib-FRIN32 indicated close genetic affinity (>98%) with *O. myriophilus*, aligning well with morphological evidence.

Phylogenetic analyses performed using maximum-likelihood methods revealed clear and robust clustering of isolates within their respective genera. Ib-CRIN68 clustered definitively with *H. bacteriophora* reference sequences, closely grouping alongside related species such as *H. indica*. Within the genus *Steinernema*, isolates Ib-IART45 and Ib-ITUC102 formed a cohesive, monophyletic clade consistent with *S. carpocapsae*, while isolates Ib-HORT and Za-SAM occupied distinct, clearly separated branches aligned respectively with *S. nepalense* and *S. feltiae*. Isolate Ib-FRIN32 demonstrated clear separation within the genus *Oscheius*, clustering closely with reference sequences of *O. myriophilus* and related species like *O. tipulae*.

The phylogenetic results verify morphological identifications and offer additional molecular clarity, establishing species identity and taxonomic relationships among the six isolates.

### Mortality Response Across Developmental Stages

The virulence of the six indigenous EPN was evaluated against four developmental stages of *S. frugiperda*: 2^nd^, 4^th^, 6^th^ instar larvae, and pupae. Mortality rates increased consistently with both nematode dose and exposure duration for all developmental stages.

Among the life stages tested, FAW 2^nd^ instar larvae were most susceptible. At 72 hours post-inoculation, all isolates achieved considerable dose-dependent mortality, with the highest mortality observed at 200 IJs/insect. Mean mortality ranged from 82.4 ± 5.6% (Ib-CRIN68) to 58.1 ± 7.2% (Ib-FRIN32) at this dose. Mortality at the lowest dose (25 IJs/insect) was modest, typically between 18% and 32% by 72 hours (Fig. 4 A-C).

**Fig. 4.**
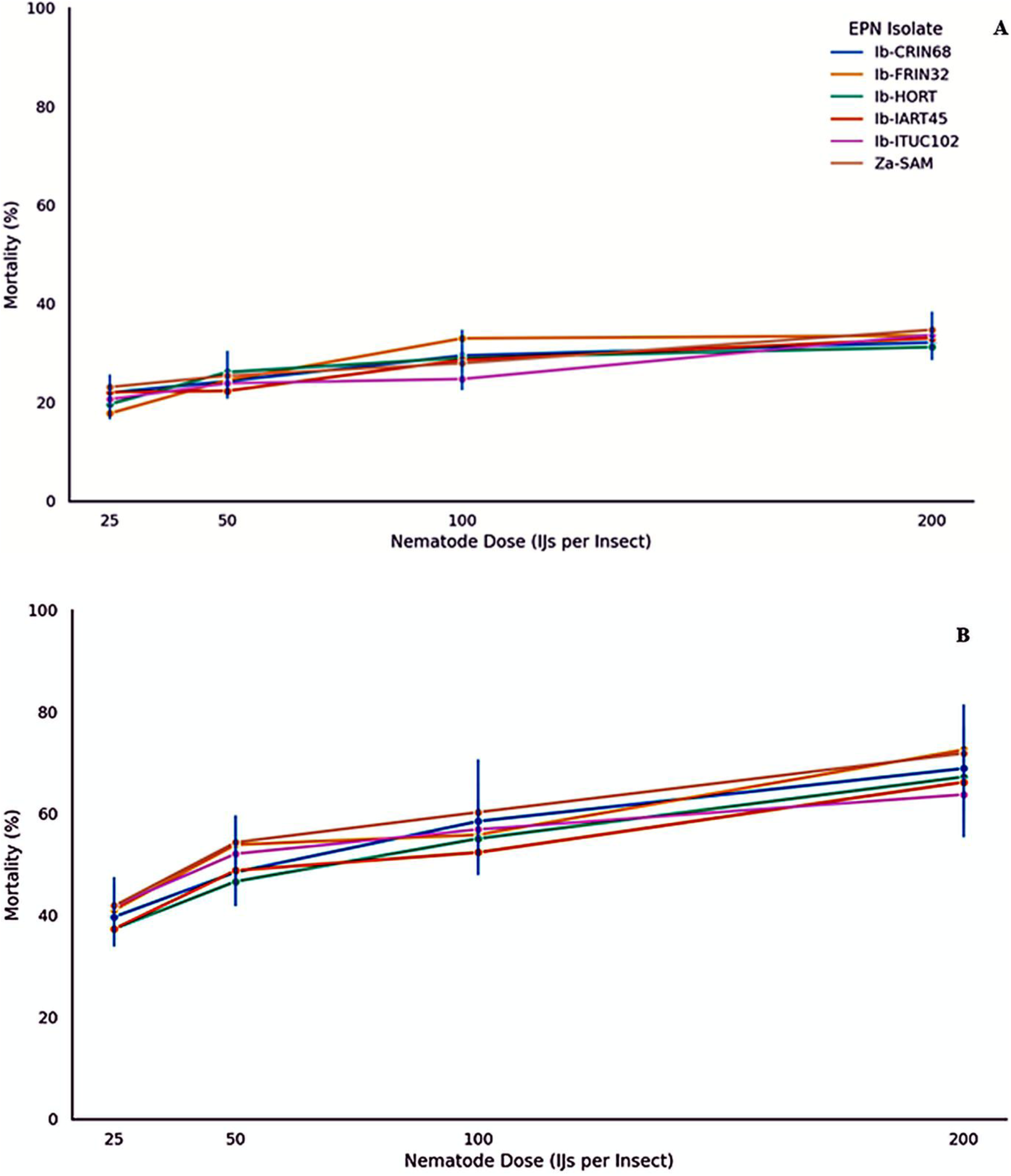
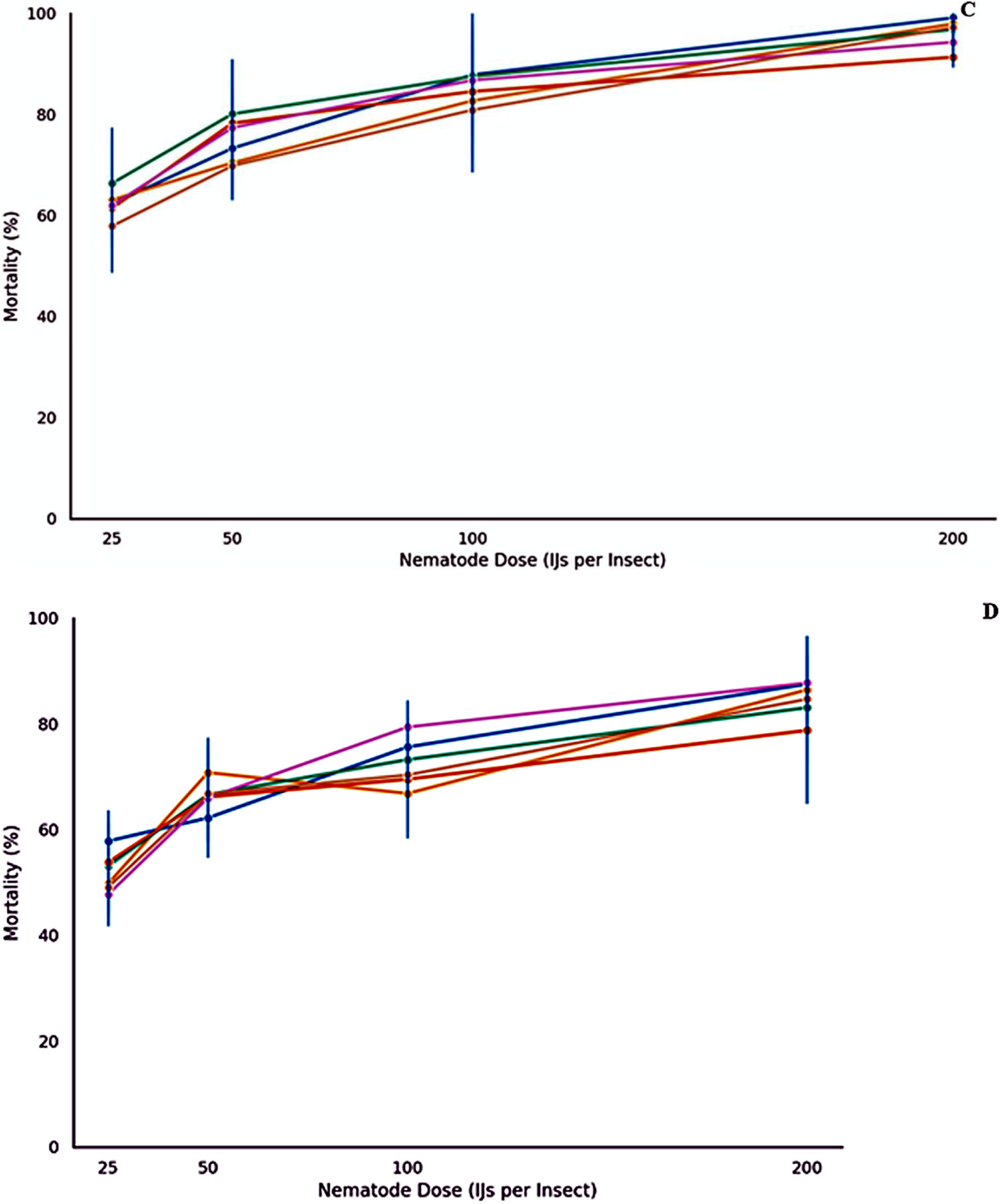
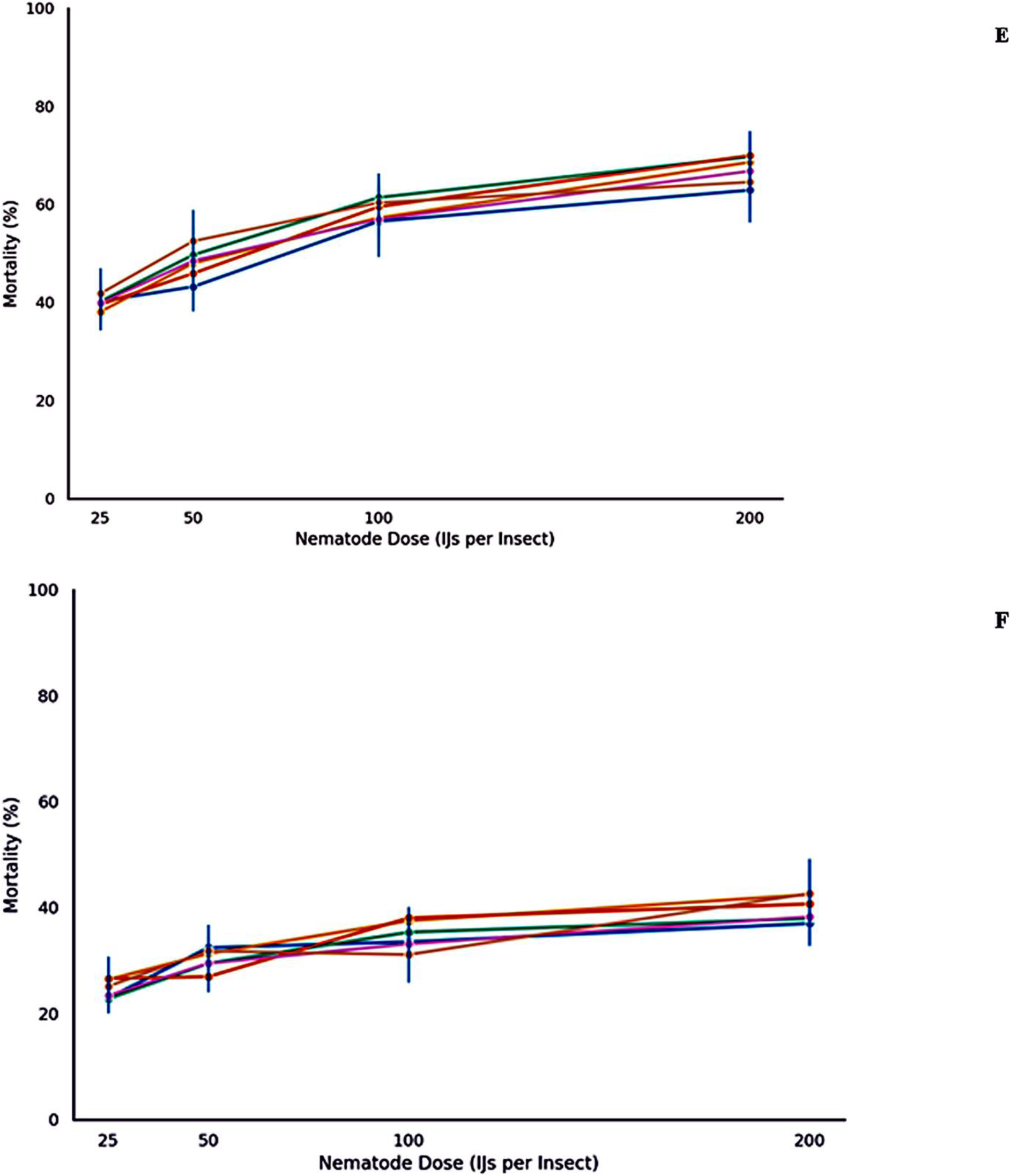
Mean mortality (% ± SE) of *Spodoptera frugiperda* larvae exposed to different dosages (25, 50, 100, and 200 infective juveniles (IJs) per insect) of six indigenous entomopathogenic nematode (EPN) isolates at different time of post-infection. 2^nd^ instar larva at 24 h, 48 h and 72h after infection (A-C); 4^th^ instar lava, 6^th^ instar larva and pupal stage at 72h after infection (D-F).

For the 4^th^ instar larvae, mean mortality ranged from 65.2 ± 8.1% to 41.7 ± 7.5% at 200 IJs/insect after 72 hours. The 6^th^ instar larvae showed lower mortality, with values between 52.5 ± 6.4% and 35.3 ± 8.7%. Pupae were the least affected, with mortality ranging from 55 ± 5.8% to 21.4 ± 6.1% across the isolates at the highest dose and latest time point.

Lethal concentration (LC₅₀) and lethal time (LT₅₀) values were calculated for the 2^nd^ instar stage based on observed dose–response trends. The isolate Ib-CRIN68 exhibited the highest potency, with an LC₅₀ of 61.8 IJs/insect and an LT₅₀ of 38.5 hours. *Steinernema carpocapsae* isolates Ib-IART45 and Ib-ITUC102 also showed favourable performance with LC₅₀ values of 72.5 and 66.0 IJs/insect, respectively. The lowest potency was recorded for the *O. myriophilus* isolate Ib-FRIN32, with an LC₅₀ of 85.3 IJs/insect and an LT₅₀ of 47.1 hours.

A three-way ANOVA confirmed that nematode isolate (F₅,₁₂₀ = 28.34, *p* < 0.001), dose (F₃,₁₂₀ = 41.76, *p* < 0.001), and time (F₂,₁₂₀ = 19.21, *p* < 0.001) significantly influenced larval mortality in 2^nd^ instars. Significant interactions were also detected between isolate and dose (F₁₅,₁₂₀ = 6.54, *p* < 0.001), and between dose and time (F₆,₁₂₀ = 5.02, *p* < 0.01).

For the 4^th^, 6^th^ instars, and pupae, one-way ANOVA performed at 200 IJs/insect and 72 hours revealed no statistically significant differences among isolates (4^th^ instar F = 0.41, *p* = 0.8296; 6^th^ instar F = 0.72, *p* = 0.6239; and pupae F = 0.95, *p* = 0.4868).

For later larval and pupal stages of *S. frugiperda*, mortality also increased in a dose-dependent manner. However, the variability in isolate performance was less pronounced compared to FAW’s 2^nd^ instar stage (Fig. 4 D-F).

At the highest dosage tested (200 IJs/insect) and 72 hours post-exposure, the efficacy of the isolates appeared to vary more notably at earlier insect developmental stages. Mortality differences among the isolates for the 2^nd^ instar larvae was not statistical significance (F = 1.93; *p* = 0.163), indicating negligible variability in isolate performance at this early stage. Also, mortality rates among isolates were statistically similar at later developmental stages, including the 4th instar (F = 0.41; *p* = 0.830), 6th instar (F = 0.72; *p* = 0.624), and pupal stage (F = 0.95; *p* = 0.487).

## Discussion

This study represents the first comprehensive attempt to isolate, morphologically and molecularly characterize, and evaluate the infectivity of native EPN strains from Nigeria against multiple life stages of FAW under laboratory conditions.

The identification of isolates Ib-CRIN68 (*H. bacteriophora*), Ib-IART45 and Ib-ITUC102 (*S. carpocapsae*), Ib-HORT (*S. nepalense*), Za-SAM (*S. feltiae*), and Ib-FRIN32 (*O. myriophilus*), corroborates previous findings and provides valuable updates to the limited EPN taxonomic records in Nigeria. Previously, (Akyazi et al., 2012) recorded the first occurrence of EPNs from Nigerian soils, identifying isolates primarily belonging to the genera *Heterorhabditis* and *Steinernema*. Our findings reaffirm the presence of these genera and expand the diversity by including the less commonly reported genus of *Oscheius*. Our identification of *O. myriophilus* in Nigerian soils is critical, as *Oscheius* spp. are emerging globally as promising biocontrol agents, particularly against dipteran and lepidopteran pests (Dichusa et al., 2021; Suan et al., 2025).

Morphological variability observed among our isolates, demonstrated through comprehensive morphometric analyses, aligns closely with recent studies conducted in similar agroecological zones worldwide (Alotaibi et al., 2022; Daramola et al., 2021). Notably, the morphological parameters employed in our study were robust enough to differentiate effectively among genera, providing essential baseline data for future comparative studies. However, relying exclusively on morphological identification can be limiting due to phenotypic plasticity of EPNs as influenced by environmental conditions, host type, and geographic isolation (Caccia, Rondan Dueñas, Del Valle, Doucet, & Lax, 2017; Guide et al., 2018; Malan, Knoetze, & Tiedt, 2016; Ngugi, Haukeland, Wachira, Mbaka, & Okoth, 2019). Thus, our integration of molecular analyses using multiple genetic markers (ITS, 12S, COI, and D2–D3 regions) considerably enhanced species resolution and reduced potential taxonomic ambiguity. This dual approach, as underscored by recent literature (Gumussoy et al., 2022; Subbotin, 2021), is crucial for correctly delineating closely related nematode species, thereby providing a reliable basis for their subsequent application in pest management programs. The comprehensive morphological and molecular characterization of EPNs isolated in this study contributes significantly to our understanding of indigenous nematode diversity in Nigerian agroecosystems and beyond.

The virulence assays conducted in this study clearly demonstrated that EPN isolates exhibit differential pathogenicity against various developmental stages of FAW. Our data revealed high susceptibility in the early instar larvae, with isolate Ib-CRIN68 (*H. bacteriophora*) achieving notably high mortality rates, consistent with prior reports by Alotaibi et al. (2022) and Guide et al. (2024). Our results also corroborate previous observations that earlier developmental stages of lepidopteran pests are typically more vulnerable to EPN attack due to their thinner cuticles, lower immune capabilities, and relatively less-developed defensive behaviours (Bai et al., 2016; Guo et al., 2023; Zhang, Mao, Liu, & Zeng, 2012). Conversely, reduced susceptibility observed in later instars and pupae of FAW in our study parallels findings from previous work that highlighted the increased physiological host defence against EPNs via thicker cuticles, enhanced immune responses, and behavioural resistance mechanisms in older FAW larvae (Acharya, Hwang, Mostafiz, Yu, & Lee, 2020; Fallet et al., 2022).

We found significant influences of nematode isolate, dosage, and exposure time on FAW mortality, particularly for early instars. Similar trends have been documented in several previous studies (Abd-Elgawad, 2019; Brusselman, Steurbaut, & Sonck, 2007; Dillon, Downes, Ward, & Griffin, 2007; Kapranas et al., 2017), emphasizing the importance of optimizing dose and exposure parameters to maximize EPN efficacy under field conditions. The absence of significant differences in virulence among isolates at later FAW developmental stages further underscores the complex interaction dynamics between host defense strategies and nematode pathogenicity (Acharya et al., 2020; Castillo, Reynolds, & Eleftherianos, 2011).

The presence of inter-isolate variation in pathogenicity among our Nigerian EPN isolates suggests potential genetic and ecological adaptations that could enhance or limit their practical effectiveness under field conditions. Thus, a more comprehensive ecological characterization, which extends beyond laboratory assays to encompass critical environmental factors such as temperature tolerance, desiccation survival, soil type preference, host-finding strategies, and compatibility with agricultural practices, is indispensable. Such detailed ecological studies have been advocated strongly in the recent literature as critical prerequisites for successful biocontrol implementation of EPNs (Koppenhöfer & Kaya, 1999; Kour, Khurma, & Brodie, 2021). Indeed, isolates exhibiting broad environmental tolerance, effective dispersal capabilities, and high persistence in the soil are more likely to provide consistent biocontrol efficacy under varying field conditions (Aryal et al., 2025; Suan et al., 2025).

Furthermore, ecological profiling of indigenous isolates is essential in tailoring biological control agents for targeted applications within specific agroecological zones. Considering Nigeria’s diverse agricultural environments, including lowland rainforest and the Guinea savannah zones represented in this study, the isolates’ ecological adaptability and specificity could significantly influence their biocontrol potential. Recent evidence from Brazilian orchards demonstrated that understanding nematode ecology substantially improved biocontrol outcomes against major pests, resulting in sustainable pest management strategies that effectively reduced chemical pesticide use (Barbosa-Negrisoli, Negrisoli, Dolinski, & Bernardi, 2010; Guide et al., 2024; Mejia-Torres & Sáenz, 2013). Possibly, our isolates could offer environmentally sustainable solutions tailored specifically to agricultural conditions in SSA where FAW represents a severe economic threat to maize production.

Hence, future research should emphasize more detailed ecological characterization of the EPN isolates identified in this study. Understanding their ecological interactions and environmental adaptability will not only optimize their biocontrol efficacy but also guide their integration into comprehensive IPM programs. Additionally, ecological characterization will help assess the feasibility of commercial-scale nematode formulations, ensuring consistency and reliability under varied field conditions.

Thus, in summary our study provides critical baseline data on indigenous Nigerian EPN isolates’ taxonomic identities and pathogenic capabilities against the invasive FAW. These findings expand the limited existing knowledge base on EPNs in SSA (Akyazi et al., 2012). Moreover, our results highlight the necessity for ecological characterization of EPNs to translate promising laboratory results into reliable field applications, thereby possibly contributing to sustainable pest management strategies in Nigeria and similar agroecological regions in SSA and beyond.

